# Reciprocal connectivity between secondary auditory cortical field and amygdala in mice

**DOI:** 10.1101/634469

**Authors:** Hiroaki Tsukano, Xubin Hou, Masao Horie, Hiroki Kitaura, Nana Nishio, Ryuichi Hishida, Kuniyuki Takahashi, Akiyoshi Kakita, Hirohide Takebayashi, Sayaka Sugiyama, Katsuei Shibuki

## Abstract

Recent studies have examined the feedback pathway from the amygdala to the auditory cortex in conjunction with the feedforward pathway from the auditory cortex to the amygdala. However, these connections have not been fully characterized. Here, to visualize the comprehensive connectivity between the auditory cortex and amygdala, we injected cholera toxin subunit b (CTB), a bidirectional tracer, into multiple subfields in the mouse auditory cortex after identifying the location of these subfields using flavoprotein fluorescence imaging. After injecting CTB into the secondary auditory field (A2), we found densely innervated CTB-positive axon terminals that were mainly located in the lateral amygdala (La), and slight innervations in other divisions such as the basal amygdala. Moreover, we found a large number of retrogradely-stained CTB-positive neurons in La after injecting CTB into A2. When injecting CTB into the primary auditory cortex (A1), a small number of CTB-positive neurons and axons were visualized in the amygdala. Finally, we found a near complete absence of connections between the other auditory cortical fields and the amygdala. These data suggest that reciprocal connections between A2 and La are main conduits for communication between the auditory cortex and amygdala in mice.

## Introduction

The auditory cortex is a center of auditory processing, and it has been found to have a functional map (Stiebler et al., 1997; Guo et al., 2011). Recent optical imaging studies have indicated that the auditory cortex of the mouse brain is composed of at least four tonotopic subfields (Issa et al., 2014) that were tentatively annotated as the dorsomedial field (DM), anterior auditory field (AAF), primary auditory cortex (A1), and secondary auditory field (A2) in a previous report (Tsukano et al., 2017). Subregions in the mouse auditory cortex may be functionally specialized (Honma et al., 2013; Baba et al., 2016; Shepard et al., 2016), and so connective patterns with other brain regions may vary according to subregion. To date, several neuronal tracing investigations have examined the afferents and efferents to/from the mouse auditory cortex. However, the injection sites have usually been set in A1. Indeed, few investigations have employed systematic injections.

Associations between auditory information and emotional meaning are biologically important because they aid in the detection of upcoming risks. The amygdala is an essential region in establishing associations between tones and emotions (Keifer et al., 2015b; Letzkus et al., 2011; Peter et al., 2012; Sah et al., 2003). Thus, full characterization of the connective pattern between the auditory cortex and amygdala is likely to produce meaningful information. To date, many studies have described the feedforward auditory–amygdala pathway that originates in the auditory cortex (Shi and Cassell, 1997; Kwon et al., 2014; Strobel et al., 2015; Manassero et al., 2018) and auditory thalamus (LeDoux and Farb, 1991; Bordi and LeDoux, 1994a,b; Kimura et al., 2003; Kwon et al., 2014; Keifer et al., 2015a). Among amygdala subdivisions, the lateral amygdala (La) is thought to be important for the convergence of auditory cues and footshock information (Romanski et al., 1993) because La receives afferents from a wide range of sensory systems including the auditory cortex (LeDoux et al., 1991; Romanski and LeDoux, 1993). A recent study characterized a feedback pathway from La to the auditory cortex in mice (Yang et al., 2016). This was consistent with the finding of older studies that used monkeys (Amaral and Price, 1984; Yukie, 2002) and rats (McDonald and Jackson, 1987). However, neuronal tracing is often conducted without clear segregation between the auditory cortical subregions. As a result, whether the whole auditory cortex or just a local area has reciprocal connections with the amygdala is unclear. This could be addressed via systematic projection mapping between the auditory cortex and the amygdala. In the current study, we visualized comprehensive connectivity between the auditory cortex and the amygdala by injecting cholera toxin subunit b (CTB), a bidirectional tracer, into four tonotopic regions of the auditory cortex, according to a map that was functionally determined using flavoprotein fluorescence imaging (Tsukano et al., 2017). We demonstrated that the A2 is specifically connected with the amygdala, which is consistent with the idea that A2 is a higher-order region in the mouse auditory cortex.

## Materials and Methods

### Animals

The experimental procedures in the present study were approved by the Committee for Animal Care at Niigata University. Data were obtained from male C57BL/6 mice (Charles River Japan, Kanagawa, Japan). The animals were housed in cages with *ad libitum* access to food pellets and water, and were kept on a 12-h light/dark cycle.

### Flavoprotein fluorescence imaging

Locations of regions in right auditory cortex were identified according to responses revealed using flavoprotein fluorescence imaging (Shibuki et al., 2003; Tsukano et al., 2016). Cortical images were recorded by a CCD camera system (AQUACOSMOS with ORCA-R2, Hamamatsu Photonics, Shizuoka, Japan) via an epifluorescence microscope (Ex, 500–550 nm; Em, 470–490 nm; M651 combined with MZ FL II, Leica Microsystems, Wetzlar, Germany). The area covered by one pixel was 20.4 × 20.4 μm. Mice were anesthetized with urethane (1.65 g/kg, i.p.; Wako, Osaka, Japan). The rectal temperature was maintained at ∼37°C. A craniotomy (∼3 × 3 mm) was performed over the right auditory cortex. The auditory cortex was activated by presentation of sound waves that were made using a LabVIEW program (National Instruments, Austin, TX). Tones were amplitude modulated (20 Hz, 100% modulation) and set to ∼60 dBSPL.

### Neural tracer injection

A glass pipette (tip diameter ∼30μm) filled with tracer solution was introduced into the center of auditory regions visualized and identified using optical imaging, to ∼450μm below the cortical surface so that tracer solutions spread to all the cortical layers. Here, we used a low-salt type cholera toxin subunit b (CTB) (#104; List Biological Laboratory, Campbell, CA) that is suitable for iontophoretic injection (Ruigrok et al., 1995). CTB was injected by carrying 70 pulses, 5 s on/ 5 s off anodal currents at an intensity of 4 μA. Partly, dual injections of Alexa Fluor 488- and 555-conjugated CTBs (0.5% in 0.1 M phosphate buffer, Thermo Fisher Scientific, Waltham, MA) were made iontophoretically by 70 pulses, each. After finishing injections, a glass pipette was slowly withdrawn. The cranial hole was covered using 2% agarose (1-B, Sigma-Aldrich, MO) and the skin was sutured. Mice were placed in a warm place for recovery, and after awaking they were reared in their home cages.

### Histology

Three days after injections, mice were anesthetized with an overdose of pentobarbital (0.3 g/kg, i.p.), and cardiac perfusion perfused transcardially with 4% paraformaldehyde. Brains were removed and immersed in 4% paraformaldehyde overnight. After incubated in 20% and 30% sucrose in 20 mM phosphate buffered saline (PBS), consecutive 40 μm-thick coronal sections were made using a sliding cryotome (REM-710, Yamato-Koki, Saitama, Japan). Every fourth slice was used for analysis.

To visualize CTB, sections were initially rinsed in 20 mM PBS and incubated in PBS containing 3% hydrogen peroxide and 0.1% Triton X-100 for 15 min at room temperature. After incubated in 20 mM PBS containing 0.1% Triton X-100 (PBST) for 60 min, slices were incubated overnight at room temperature with goat anti-CTB antibody (List Biological Laboratories) diluted to 1:30,000 with 20 mM PBS containing 0.5% skim milk. The next day, slices were rinsed in 20 mM PBS, and incubated at room temperature for 2 h with HRP-conjugated rabbit anti-goat IgG antibody (MBL, Nagoya, Japan) diluted to 1:200 using 20 mM PBS containing 0.5% skim milk. Sections were rinsed in 20 mM PBS, and the immunoreactions were visualized in a Tris-HCl buffer containing 0.05% diaminobenzidine tetrahydrochloride and 0.003% hydrogen peroxide for 5 min at room temperature. After visualization, slices were Nissl-stained using 0.1% cresyl violet (Chroma Gesellschaft, Kongen, Germany), and they were dehydrated in ethanol, cleared in xylene, and cover-slipped using the covering reagent Bioleit (Okenshoji, Tokyo, Japan). When coverslipping sections with fluorescent CTB, Fluoromount (Cosmo Bio, Tokyo, Japan) was used as a covering reagent instead.

Borders of subdivisions in the amygdala were drawn according to the mouse brain atlas (Paxinos and Franklin, 2012) and Nissl staining patterns. Histological images were obtained using a CCD camera (DP80; Olympus, Tokyo, Japan) via a stereoscopic microscope (Eclipse Ni, Nikon, Tokyo, Japan) using white light or emission filters (515–555 nm for greed and 600–660 nm for red).

### Statistics

The Mann-Whitney U test or Wilcoxon signed-rank test was used to evaluate differences between unpaired or paired data from the two groups, respectively. If multiple comparisons were in need, p-values were corrected using the Bonferroni correction. The Dunnett’s test was used to evaluate multiple data against reference data. The chi-squared test was used to evaluate differences between two distributions. All tests were conducted as a two-sided test using MATLAB programs (Mathworks, St. Louise, MO) or SPSS (IBM, Armonk, NY). All data in graphs are presented as mean ± S.E.M.

## Results

### Auditory cortical region-specific communications with the amygdala

The mouse auditory cortex includes at least four tonotopic regions (Issa et al., 2014; Tsukano et al., 2017), which we refer to here as DM, AAF, A1, and A2 (Fig. 1A,B). After identifying the precise location of AAF, A1, A2, and DM using flavoprotein fluorescence imaging, we injected CTB into the center of the responsive region to examine connective patterns with the amygdala. Deposits of CTB localized within the injected regions were confirmed after preparing brain slice sections (Fig. 1C).

**Figure 1.**
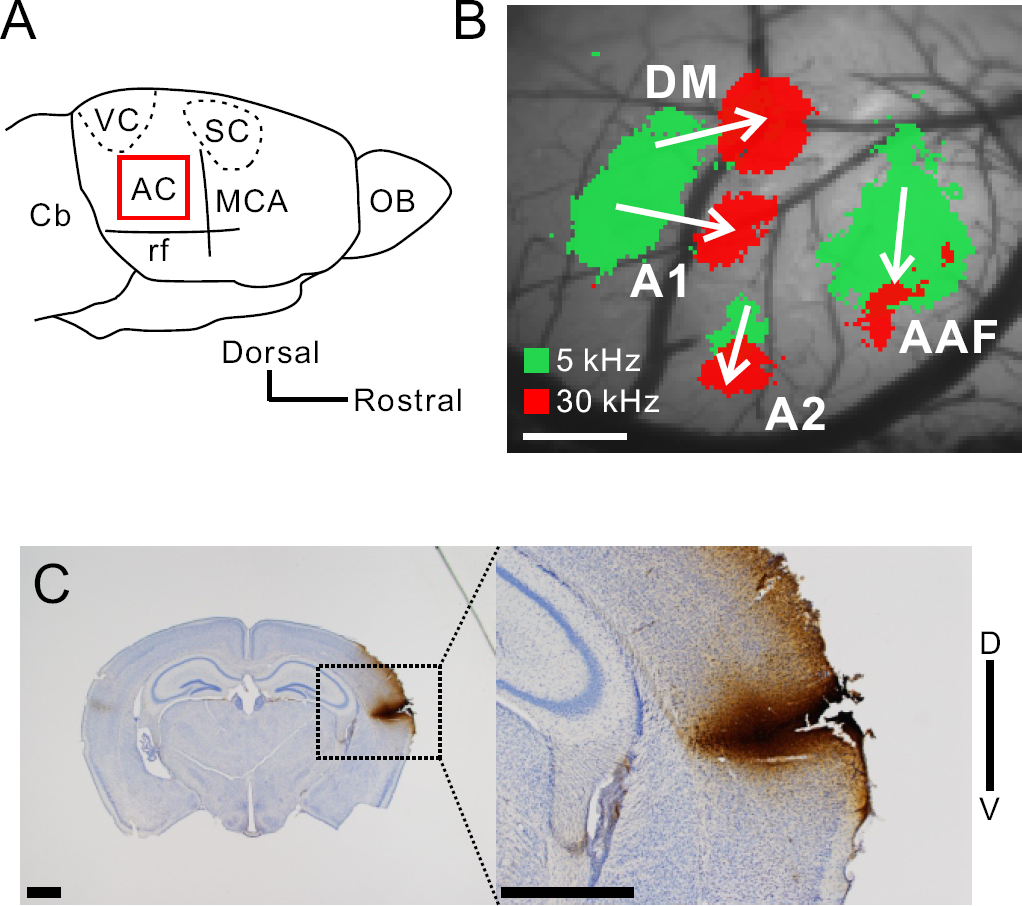
Injection of neural tracers into auditory regions. **A**, An illustration of the right auditory cortex in mice. AC, auditory cortex; Cb, cerebellum; MCA, medial cerebral artery; OB, olfactory bulb; rf, rhinal fissure; SC, somatosensory cortex; VC, visual cortex. **B**, Identification of auditory cortical regions with the guidance of tonal responses via flavoprotein fluorescence imaging. Scale bar, 500 μm. AAF, anterior auditory field; A1, primary auditory cortex; A2, secondary auditory field; DM dorsomedial field. **C**, An example showing CTB injection sites after preparation of coronal slices. Scale bar, 1 mm.

We examined the distribution of CTB-positive axon terminals projected from the auditory cortex in the amygdala. The images shown in Figure 2 were obtained by injecting CTB into A2. We found that densely distributed CTB-positive axon terminals were located mainly in La, and identified slight innervations in other divisions such as the central amygdala (CeA) and basal amygdala (BA), including the anterior basolateral amygdala (BLA), posterior basolateral amygdala (BLP), anterior basomedial amygdala (BMA), and posterior basomedial amygdala (BMP) (Fig. 2). As for the three intercalated nuclei, which are a cluster of GABAergic neurons, the two dorsal nuclei the sides of La included some CTB-positive terminals. This is consistent with a previous study (Strobel et al., 2015). The most well-known cluster, which was located ventrally, included no CTB-positive axon terminals. These data suggest that La mainly receives input from the auditory cortex, and this result is consistent with the well-known view that La is an entrance in the amygdala for sensory input from sensory cortices (Sah et al., 2003). Moreover, we found many CTB-positive somata, located mainly in La, after injecting CTB into A2 (Fig. 2). We quantitatively analyzed the number of CTB-positive somata in each amygdala division (Fig. 4A). Numerous CTB-positive neurons were distributed in La, and we found a small number of neurons in the basolateral and basomedial amygdala. Importantly, we found no CTB-positive somata in CeA (Fig. 4A), despite the proposed role of CeA as the output center of the amygdala (Keifer et al., 2015b). These data confirm that A2 projects mainly to La and receives direct feedback projections from La but not from CeA.

**Figure 2.**
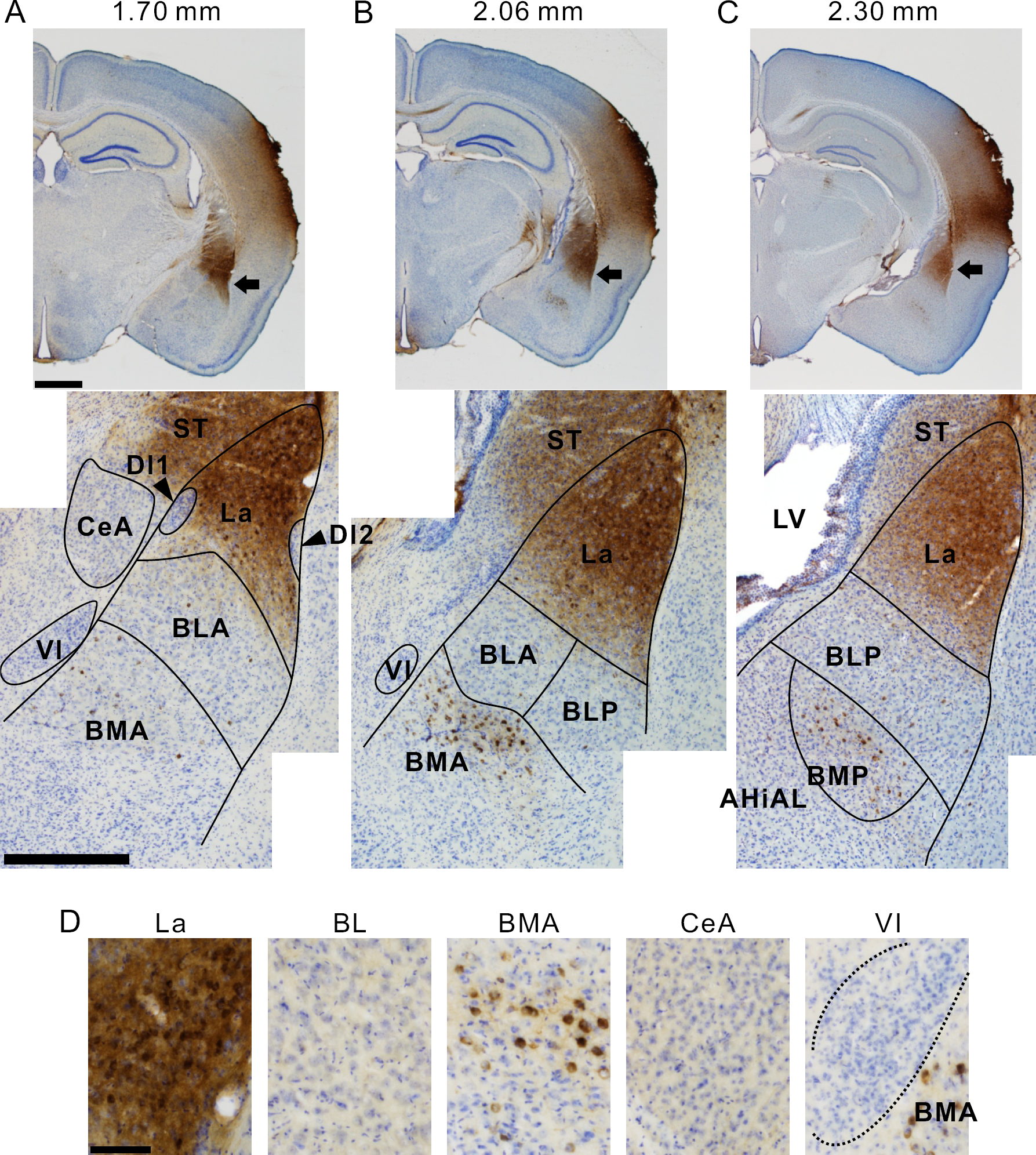
CTB-positive neurons and axon terminals in communication with the auditory cortex. **A**, Low and middle magnification images of the amygdala 1.70 mm posterior to the bregma. Arrows indicate the CTB-positive amygdala neurons. Scale bar, 1 mm (top) and 500 μm (bottom). **B**, A pair of images taken 2.06 mm posterior to the bregma. **C**, A pair of images taken 2.30 mm posterior to the bregma. The arrow head indicates CTB-positive neurons in the higher-order visual thalamus–lateral posterior thalamus (LP). **D**, High magnification images of subdivisions of the amygdala. Scale bar, 100 μm. AHiAL, anterolateral amygdalohippocampal area; BL, basolateral amygdala; BLA, anterior BL; BLP, posterior BL; BMA, anterior basomedial amygdala; BMP, posterior basomedial amygdala; CeA, central amygdala; DI, dorsal intercalated nucleus; La, lateral amygdala; LV, lateral ventricle; ST, striatum; VI, ventral intercalated nucleus.

**Figure 3.**
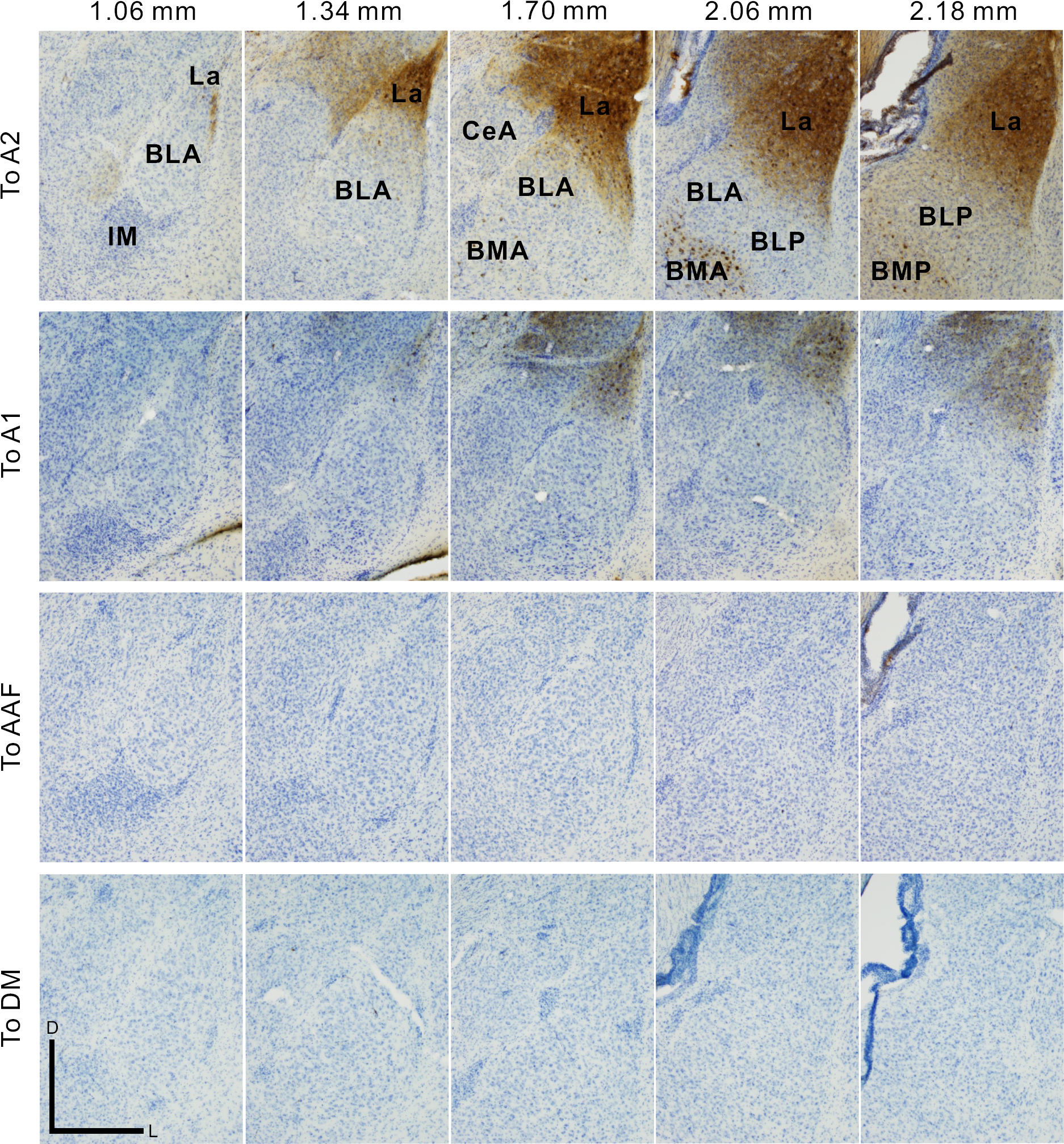
A2- and A1-specific communications with the amygdala. The panel shows the amygdala in the coronal slices after CTB was injected into A2, A1, AAF, or DM in the auditory cortex. IM, main intercalated nucleus. Scale bar, 500 μm.

**Figure 4.**
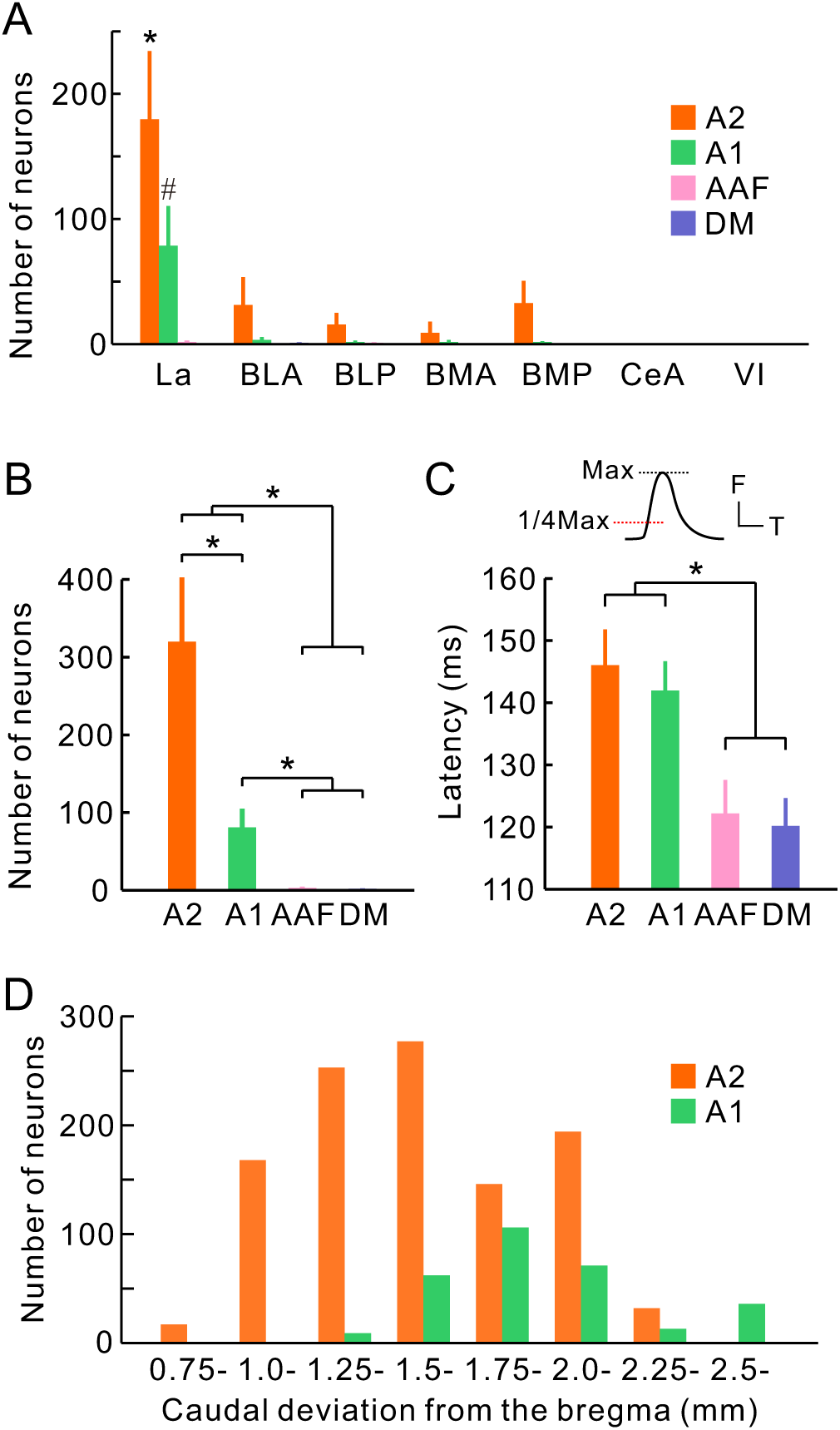
Quantitative analysis of connections between the auditory cortex and the amygdala. **A**, Quantitative analysis of the distribution of CTB-positive neurons in the amygdala. We found more CTB-positive neurons in La compared with the other subdivisions (*p < 0.001, #p<0.0001, Dunnett’s test). Data were obtained from five, four, three, and four mice for A2, A1, AAF, and DM, respectively. **B**, Number of CTB-positive neurons in the amygdala by injection site. *p < 0.05, Wilcoxon signed-rank test after the Bonferroni correction. **C**, Hierarchical relationships between auditory cortical regions, represented via differences in latencies. Latency represents the time that is required to reach 1/4 maximum of the peak response. The graphs were obtained from the same dataset published in Tsukano et al., 2015 (n = 18 mice each; *p < 0.01, Wilcoxon signed-rank test after the Bonferroni correction). F, fluorescence; Max, maximum; T, time. **D**, Distribution of CTB-positive neurons projecting to A2 and A1 in the rostrocaudal axis. Two distributions were significantly different (p < 10^-6^, chi-squared test). The same sets of brain slices were used for (A), (B), and (D).

We remarked that the density of CTB-positive axon terminals and the number of CTB-positive neurons in the amygdala differed by injection site in the auditory cortex. As described above, A2 has strong connections with La (Figs. 3 and 4A,B). After injecting CTB into A1, we found one third the number of CTB-positive neurons compared with that observed after injecting CTB into A2 (Figs. 3 and 4A,B). The amount of CTB-positive axons was proportional to the number of CTB-positive neurons, indicating that A1 also had few reciprocal connections with La. Furthermore, we did not find CTB-positive axon terminals or CTB-positive neurons inside the amygdala after injecting CTB into AAF and DM (Figs. 3 and 4A,B). Interestingly, the numbers of CTB-positive neurons in the amygdala increased in proportion with the hierarchical order of the injected regions. This was reflected in the tonal response latencies, which were measured as quarter-maximal latencies of sound-induced flavoprotein fluorescence responses (Fig. 4C). These data indicate that A2 is reciprocally connected with La and that it is a main conduit for communication with the amygdala, while A1 has connections that are similar but much weaker in strength.

### Two independent streams between the auditory cortex and La

Many deep brain parts have initially been considered to be single structures and are then found to actually contain several substructures that can be classified according to relationships with cortical targets (Horie et al., 2013; Tohmi et al., 2014). To determine whether the origins of the La–A2 and La–A1 pathways are the spatially similar or different inside La, we evaluated the locations of La neurons projecting to A2 and A1. We found that amygdala neurons projecting to A2 were widely distributed rostrocaudally, while those projecting to A1 were comparatively localized in a caudal region (Fig. 4D). However, the distributions of amygdala neurons projecting to A2 and A1 were largely overlapped rostrocaudally. Therefore, we conducted double-injection of two fluorescence CTBs into A2 (green) and A1 (red) to investigate whether La neurons projecting to A1 and A2 were intermingled or topographically-segregated in the coronal view (Fig. 5A). We found that the two neuronal groups were distributed in a salt-and-pepper fashion and exhibited no clear spatial separation (Fig. 5B,C). However, most neurons were labeled with either tracer, and only 3.9% neurons were double-labeled (Fig. 5D). These data indicate that, although La is a single structure containing neurons with different cortical targets that are intermingled, the La–A2 and La–A1 pathways are mutually exclusive and mediated by discrete populations of amygdala neurons.

**Figure 5.**
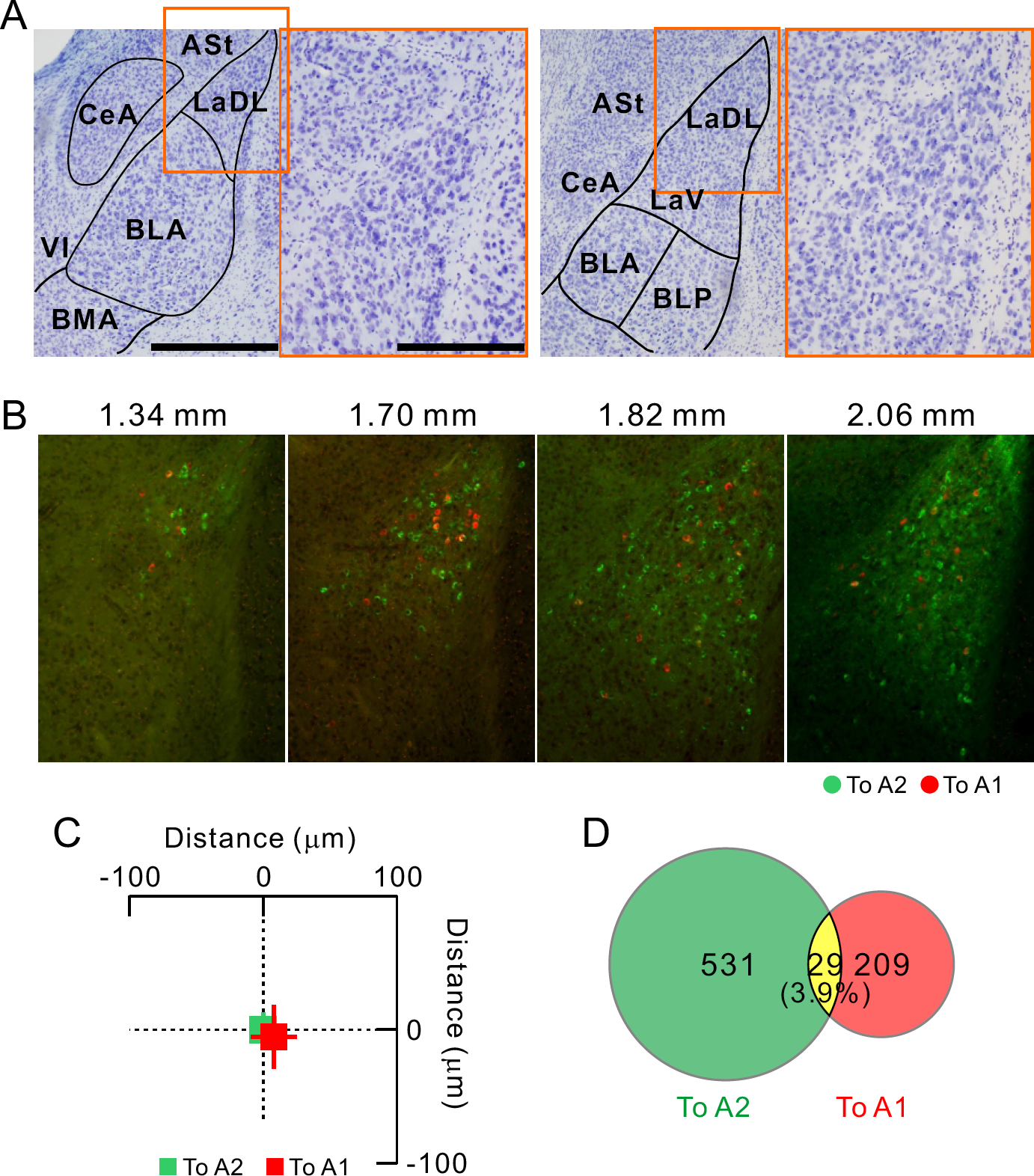
Mutually exclusive La neurons projecting to A1 and A2. **A**, Coronal views of LaDL 1.58 mm (left) and 1.94 mm (right) posterior to the bregma. The high magnification images show the regions enclosed by the orange rectangles in the low magnification images. Scale bars, 500 μm in the high magnification images and 250 μm in the low magnification images. ASL, amygdalostriatal transition; LaDL, dorsolateral part of the La; LaV, ventral part of La. **B**, Distribution of La neurons projecting to A2 (green) and A1 (red). Scale bar, 250 μm. **C**, No spatial topography in neurons projecting to A1 and A2. The averaged coordinates of neurons projecting to A2 were set as the origin. The slices on which both neuronal groups were visualized were used for analysis. We found no significant topographic shifts in the horizontal (p > 0.8; Mann–Whitney U-test) or vertical axes (p > 0.4; Mann–Whitney U-test). Data were obtained from two mice; to A2, n = 282 neurons; to A1, n = 102 neurons. **D**, Proportion of neurons projecting to the A2 and/or A1. Data were obtained from four mice; to A2, n = 531 neurons; to A1, n = 209 neurons.

The auditory cortex has tonotopic organization based on tonal frequencies. It is already known that the auditory cortex is connected to several deep brain structures in a frequency-related, topographical fashion, as indicated by auditory cortico-striatum projections (Xiong et al., 2015). To examine whether La and A2 are connected topographically, we injected two fluorescent CTBs into low and high frequency areas of A2 (Fig. 6). Although the distribution of the two neuronal groups was wider and more intermingled, the averaged locations of the two neuron types were significantly shifted in the dorsoventral direction on all rostrocaudal levels (Fig. 6B,C). These data suggest that the feedback projections from the amygdala to the auditory cortex have a frequency-related, topographic organization.

**Figure 6.**
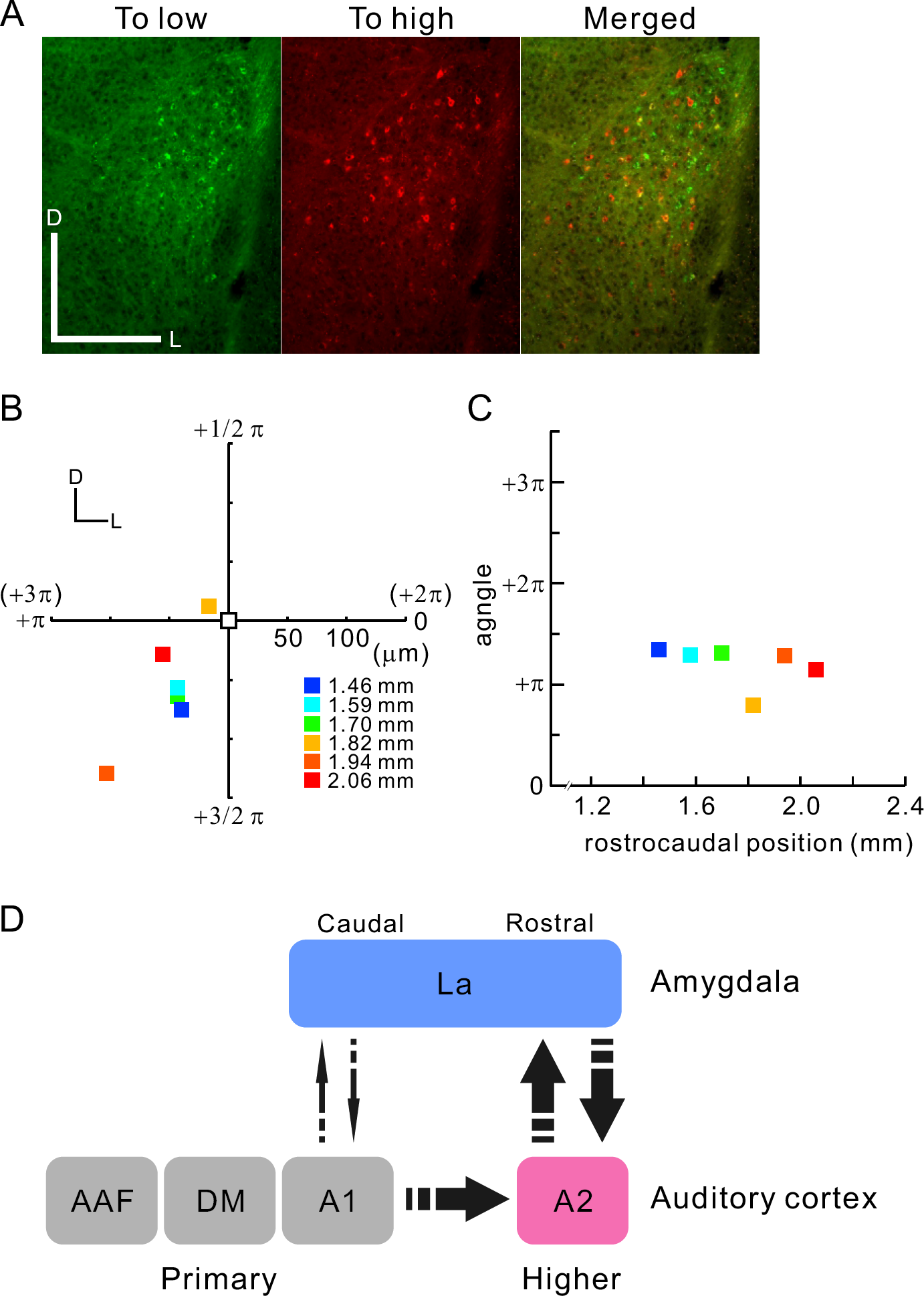
Frequency-related topography in projections between La and A2. **A**, Example slices showing La neurons projecting to low and high frequency areas of A2. fCTB-positive somata are plotted in the right panels in reference to real images shown on the left. Scale bar, 250 μm. **B**, Relative locations of La neurons projecting to low and high frequency areas of A2. The averaged coordinates of neurons projecting to low frequency area of A2 are shown in white and set as the origin, and those projecting to high frequency areas are shown in the same Cartesian coordinate system in color. Colors signify rostrocaudal levels of coronal slices in reference to the bregma. Azimuths of color plots are also shown in radians in the polar coordinate system, in which one round corresponds to 2π radians starting from the positive part of the horizontal axis. Data were obtained from four mice. Numbers of neurons were 59 at 1.46 mm, 48 at 1.58 mm, 187 at 1.70 mm, 93 at 1.82 mm, 15 at 1.94 mm, and 67 at 2.06 mm posterior to the bregma. **C**, Correlations between rostrocaudal location and azimuths in (B), r = 0.77, p > 0.07, Spearman’s test. **D**, Summary of the current results showing the connectivity between the auditory cortex and amygdala in the mouse brain.

## Discussion

The amygdala complex is important for associating affective information with sensory information. On the anatomical ground that La receives afferents from a wide range of locations in the thalamus and sensory cortex, auditory cues and footshock information converge in La (Romanski et al., 1993) and are associated with one another based on synaptic plasticity (Kessels and Malinow, 2009; Park et al., 2016). Therefore, La is considered to be the center to the establishment of fear conditioning (Sah et al., 2003). The associated information in La is then sent to CeA, which is indispensable for expressing freezing behaviors (Killcross et al., 1997). The output from CeA is then sent to a wide range of brain regions including the bed nucleus, hippocampus, midbrain, pons, and medulla oblongata (Sah et al., 2003; Keifer et al., 2015b). This general scheme illustrates La as the input center and CeA as the output center of the amygdala. However, sensory cortices, including the auditory cortex, have not been found to be output targets of CeA. Indeed, old tracing studies had already revealed that the auditory cortex receives dense projections from the lateral parts of the amygdala in rats (McDonald and Jackson, 1987) and primates (Yukie, 2002). Both our findings and those of a recent study (Yang et al., 2016) confirm the presence of direct projections from La to the auditory cortex in mice, as well as the absence of projections from CeA to the auditory cortex. A similar scheme is also applicable to the visual system. The postrhinal area (POR) is a higher order visual cortical areas that receives direct feedback projections from La but not from CeA in mice, and is modulated by La during processing of motivationally relevant sensory cues (Burgess et al., 2016). Therefore, it is reasonable to classify La, and not the CeA, as the main center in the amygdala that outputs information to the sensory cortex.

The results of the current study indicate that A2 specifically outputs and inputs signals to/from La. Although previous studies have reported evidence of a direct feedback pathway from La to the auditory cortex both in mice (Yang et al., 2016) and rats (McDonald and Jackson, 1987), whether the entire auditory cortex is uniformly innervated by La has not been determined. As for the feedforward pathway from the auditory cortex to the amygdala, a consensus has not been made regarding whether the projections originate throughout the entire auditory cortex or are localized. Interestingly, one tracing study suggested that the ventral parts in the rat auditory cortex give rise to projections towards the amygdala (Shi and Cassell, 1997). However, the stereotaxic identification of the auditory cortex in this study was based on the classical regional annotation of Te1, Te2, and Te3 (Zilles 1990), which is quite different from the modern classification using AAF, A1, and A2. In the current study, the combination of neuronal tracing with prior functional identification of the auditory subregions using optical imaging enabled us to reveal the precise projectional topography based on the modern map of the auditory cortex. Thus, this study is the first to show that both the feedforward and feedback pathways between the auditory cortex and amygdala do not uniformly originate or terminate across the entire auditory cortex or amygdala in mice, and that A2 and La are main conduits for communication between the auditory cortex and amygdala.

Overall, our findings endorse the notion that A2 is a higher-order auditory region in the mouse auditory cortex. The auditory subregions of DM, AAF, A1, and A2 have been found to be tonotopic (Tsukano et al., 2017), and all of these fields receive substantial projections from the lemniscal auditory thalamus of the ventral division of the medial geniculate body (MGv) (Tsukano et al., 2017; Ohga et al., 2018). Because the general concept explains that cortical areas that receive afferents from MGv are primary auditory areas, a question arises regarding whether these fields, including A2, are all primary areas in the mouse cortex. However, the results of the current study clearly shows the presence of dense connections between A2 and the amygdala, while other auditory fields have slight or no connections with the amygdala. Moreover, previous studies have reported that A2 neurons have long latencies (Kubota et al., 2008; Guo et al., 2012; Tsukano et al., 2015), wide bandwidth (Issa et al., 2014; Ohga et al., 2018), and a less-ordered tonotopic arrangement (Issa et al., 2014; Ohga et al., 2018). These data confirm that mouse A2 has properties of a higher-order region, and also support the idea that auditory cortical subregions are functionally specialized in mice (Honma et al., 2013; Baba et al., 2016; Issa et al., 2017). Given that primary-like regions have few feedforward or feedback connections with the amygdala, they may work to analyze auditory information, as previously proposed. Although our data showed a relatively long latency in the region labeled A1, the connectivity with the amygdala was faint, indicating that A1 may function as a primary-like region. In contrast, A2 may work in cooperation with the amygdala to associate affective information with auditory information after it receives auditory information from primary fields via the corticocortical pathways (Lee and Winer 2008; Covic and Sherman 2011). A recent study using rats suggested that value-coding neurons exist in parts that correspond to those located near mouse A2 (Grosso et al., 2015; Concina et al., 2019), which may be evidence of the La–A2 feedback pathways. It is possible that A2 works to provide biological meaning for emotion-arousing sounds such as conspecific courtship songs and predatory calls, and thus plays a more important role in signal processing than simple sound analysis.

## Author contributions

H.Ts, X.H. M.H., S.S., and K.S. conceptualized and designed the study. H.Ts, X.H., M.H., H.K. and R.H. conducted histological experiments. H.Ts, X.H., M.H., H.K., and N.N. analyzed data. N.N., K.T., K.A., S.S., and H.Ta provided critical ideas and comments regarding the research. H.Ts, X.H., and M.H. prepared the figures. H.Ts wrote the manuscript. H.Ts, X.H., S.S., and K.S. revised the manuscript. H.Ts obtained the funding. All the authors approved the publication of the manuscript.

### Acknowledgements

This work was supported by JSPS KAKENHI Grant No. 17K07051 (to H. Tsukano) and No. 26830008 (to H. Tsukano), a grant for the Promotion of Medical Science and Medical Care No. 15KI149 from the Ichiro Kanehara Foundation (to H. Tsukano), and a grant for Basic Science Research Projects No. 140254 from the Sumitomo Foundation (to H. Tsukano). We thank S. Maruyama for technical assistance and M. Isogai for conducting animal breeding and maintenance. We thank S. Koke, MFA, from Edanz Group (www.edanzediting.com/ac) for editing a draft of this manuscript.

